# Obtaining high-resolution cryo-EM structures using a common LaB6, 120-keV electron microscope equipped with a sub 200-keV optimised direct electron detector

**DOI:** 10.1101/2024.05.26.595910

**Authors:** Hariprasad Venugopal, Jesse Mobbs, Cyntia Taveneau, Daniel R. Fox, Ziva Vuckovic, Gavin Knott, Rhys Grinter, David Thal, Stephen Mick, Cory Czarnik, Georg Ramm

## Abstract

Cryo-electron microscopy (Cryo-EM) single particle analysis (SPA) has become a major structural biology technique in recent years. High-resolution cryo-EM typically requires higher voltage cryo-TEMs with coherent FEG sources, stable columns, autoloader systems and direct electron detectors. These setups are specialised for Cryo-EM work and are expensive to establish and maintain. More recently the concept of using 100-keV cryo-TEMs has been introduced as a way to make cryo-EM more affordable and hence accessible to a larger group of researchers. So far, the implementation of these 100-keV cryo-TEMs have relied on specialised microscopes with FEG sources as well as more stable optics than usually present on the common 120-keV TEMs. We here explored whether a standard 120-keV TEM, commonly available at many laboratories worldwide, can be upgraded with a direct electron detector and its suitability for high-resolution cryo-EM using a standard side entry cryo-holder. Using this imaging configuration, we were successful in achieving a 2.65Å reconstruction for standard apoferritin. We were also able to resolve a more challenging small 64kDa protein haemoglobin to 4.33Å. Furthermore, we were able to solve an asymmetric 153 kDa membrane protein GPCR (M4 muscarinic acid receptor) to a resolution of 4.4Å. Importantly, all these results were achieved using a standard automated data collection routine implemented through SerialEM, making it feasible to collect large cryo-EM data sets with a side entry cryo-holder. These results showcase a potentially widely accessible solution to obtaining interpretable cryo-EM structures. Furthermore, we envisage that this imaging configuration gives an option for many EM facilities and laboratories to set up a high-quality cryo-EM SPA sample screening capability without the need to procure costly specialised Cryo-TEMs. This could help to considerably lower the economic entry barrier for cryo-EM SPA and contribute to the “democratisation” of cryo-EM.

## Introduction

In the past decade, cryo-electron microscopy (cryo-EM) single particle analysis (SPA) has become a major technique for structural biology [1], excelling in areas where protein crystallography has been limited [2]. The technical infrastructure needed for this endeavour involves specialised cryo-EM microscopes with high coherence electron field emission gun (FEG), high operational vacuum for the stability of the FEG, improved columns, constant powered lenses and specialised cryo-stages. Together these provide the optical and mechanical stability required for obtaining a typical high-resolution SPA dataset during 12-24 hrs automated data collection. Given that improvements in electron source, column and optics were already available for the materials sciences, the development of direct electron detectors is likely the most critical innovation in aiding low-dose imaging and achieving the “resolution revolution” in cryo-EM [3]. Pushing the signal to noise level beyond film, originally used in early cryo-EM studies, these detectors were in first instance designed to yield optimal detective quantum efficiency (DQE) for imaging 200-300 keV electrons [4–6]. These detectors also made it possible to fractionate the total dose required for a typical exposure (50-60 *e^−^ Å^−2^*) into subframes. This, combined with the advancements in motion correction algorithm[7] facilitated the efficient correction of beam-induced sample drift, as well as residual stage drift in the individual exposures. This made it possible to routinely achieve resolutions below 4Å for biological specimens[3]. Additionally, many notable innovations in recent years have boosted the achievable resolution in Cryo-EM. This includes the incorporation of cold FEGs for Cryo-EM[8], the development of a stable energy filter to enable zero loss filtering using a narrow slit width of 10eV or below [9], and the improvements in minimising detector readout noise namely, correlated double sampling (CDS) mode in Gatan’s K3 detector [10] and multiframe CDS implementation in Falcon 3-4 detectors (Thermo Fischer Scientific). Individually or combined, these innovations have made it possible to solve novel biological structures to sub 2Å resolution [11–15].

For most single particle projects, the total imaging time required for its successful completion can be divided into screening and optimising sample preparation conditions and the final high-resolution imaging. Screening usually involves assessing grid preparation by imaging multiple grids in order to identify the conditions that result in optimal particle distribution and quality of ice. During the screening, a limited dataset from 2-3 grid-squares with varying ice thickness is assessed for best particle distribution from each grid. This exercise is also useful to assess potential pitfalls in cryo-EM grid preparation such as preferential orientation, air-water interface or to understand problems arising from biochemistry like intactness of biological complexes and flexibility [16–18]. Ideal cryo-EM samples would typically yield class averages showing multiple orientations as well as secondary structure level details at ∼6-8 Å resolution. A 3D reconstruction from such a dataset could potentially yield 3-6 Å resolution, enough to screen for complex formation, antibody or fab binding and even for the presence of peptides or small molecules (at ∼3-4 Å resolution). Once the grid making conditions have been established from limited processing, the same grid or replicates can be sent for imaging at state-of-the-art 300-keV microscopes such as the TFS Titan Krios or the JEOL cryo-ARM300 for the collection of large highest quality data sets [19].

The screening process can easily amount for the largest proportion of imaging time required for the successful completion of a project, especially for more challenging samples. It should be noted that higher quality data sets for screening purposes require relatively expensive cryo-TEMs typically having a 200-keV FEG source, constant power lenses, direct electron detectors and often autoloader stages. In this context, lately, potential benefits have been highlighted for performing cryo-EM at 100-keV [20] and the application for high-resolution structure determination was shown using a modified DECTRIS EIGER X 0.5 mega pixel detector on a TF20 operated at 100 keV [21]. Since then, a new dedicated 100-keV microscope was developed, featuring a newly designed 1 mega pixel dedicated 100-keV detector (DECTRIS SINGLA), a low Cc objective lens and a Schottky FEG (S-FEG) electron source with compact power supply. The design of this microscope especially in its non-requirement of the greenhouse gas, sulphur hexafluoride (SF6), for insulation, considerably reduced the cost, complexity and environmental impact [22]. This system is not yet commercially available, but these studies show the promising future of cheaper microscopy solutions enabling democratisation of the field [16, 22, 23]. Taking a lead from these studies, commercial manufactures have made strides into developing sub 200-keV detectors. This includes the Alpine direct electron detector by GATAN, Falcon-C by Thermo Fisher Scientific, and Quantum C100 from Quantum detectors. Recently, a newly developed microscope by Thermo Fischer Scientific called Tundra has been shown to be an effective tool for sample screening using 100-keV imaging with the Falcon-C detector [23]. However, this imaging configuration is still a relatively expensive microscope due to its X-FEG source, higher quality column with constant power objective lens and semi-automated sample loader. One of the main bottlenecks for wide adoption of single particle cryo-EM hence is the high cost of these imaging capabilities as well as the service contracts to maintain them.

In the context of commercially available sub 200-keV optimised direct electron detectors, we looked for further opportunities for making high-resolution cryo-EM more affordable. We focused especially on the sample screening aspect of Cryo-EM SPA which requires the ability to reach a resolution of 3-4 Å in the best-case scenario. However so far, these detectors have only been tested on costlier FEG source instruments, but not yet on standard 120-keV TEMs with a common LaB6 thermionic source. There are few published cases where LaB6 filaments were used to obtain 6-10 Å resolution structures imaged at 400keV using a Jeol JEM-4000 [24–26], but these studies predate the introduction of direct electron detectors and were imaged on film. The major inherent factor limiting resolution is the spatial and temporal coherence of the LaB6 source. The energy spread for the LaB6 source is typically in the range of 1.5-2eV as opposed to <0.8-0.9 eV for Schottky FEG [27] XFEG 0.7eV and <0.3 eV for cold FEG[9]. However, even taking these limitations into account, the line and point resolution of a typical 120-keV LaB6 TEMs is in the range of 2.0Å and 3.6Å respectively. In addition, practical limitations such as the substantial sample drift of side entry cryo transfer holders, insufficient environmental isolation and stability in terms of vibration and acoustical noise can further limit the performance. Since dose fractionated data from direct electron detector can be used to mitigate problems with stage drift and beam-induced motion, we envisaged that the addition of such a detector on a traditional 120-keV LaB6 microscope can hugely boost its performance. For the present study, a standard Tecnai 120-keV LaB6 G2 spirit TWIN microscope (Themofischer Scientific) was retrofitted with Gatan’s sub 200-keV optimised direct electron detector Alpine. At 100 keV, the Alpine detector has been shown to have 4-fold better DQE than Gatan’s K3 at the Nyquist frequency [28]. We characterise this imaging configuration for its utility to derive high-resolution structural data. In addition to well behaved samples, we show that challenging proteins as well sub 100kDa proteins can be resolved to higher resolution using this configuration. This could be a more affordable and widely available option for high quality cryo-EM SPA for both screening purposes as well as structure determination.

## Results

### A direct electron detector improves imaging on a 120-keV LaB6 TEM

To test the impact of a direct electron detector on the performance of a standard 120-keV cryo-capable TEM, we used a Tecnai G2 Spirit with a TWIN objective lens (non-constant powered lens) with a Cs of 2.2 mm and Cc of 2.2 mm. The electron source used is a standard LaB6 thermionic source with 15µm flat tip and 90° cone angle (DENKA). As these instruments are not typically used for high-resolution data collection, the housing conditions for such microscope are usually not optimal. With low DQE CCD cameras traditionally fitted with these microscopes combined with poor vibration isolation and poor stage drift, observing features close to line resolution of 2.0 Å tends to be difficult but not impossible.

After tuning the microscope for 120keV, a cross grating sample mounted on room temperature holder was imaged to assess the optical performance of the microscope in parallel illumination conditions. Once the stage drift settled it was possible to observe 2.35 Å lattice signal as observed in selected area FFT from an HRTEM image captured using a BM-Eagle CCD detector (figure 1D). The 4 s exposure did result in capturing some residual drift as seen in a directional loss of information in the power spectrum (figure 1B). This camera was unmounted and was replaced with the commercially available GATAN sub 200 keV optimised Alpine detector (supplementary figure 1 A-C). As it is a direct electron detector, imaging with alpine detector allowed us to save images as dose fractionated movies which allows for post-acquisition drift correction. Optical alignments of the microscope were checked and a longer exposure of 8 seconds was chosen to capture stage drift and environmental vibrations that might affect data collection during typical usage. The resultant exposure showed severe motion blurring due to drift (figure 2A), which is also visible in the associated power spectrum (figure 2B). Digital micrograph’s built in motion correction tools were used to analyse and further correct the drift post acquisition. The drift analysis showed that there was ∼1.6 nm cumulative drift in Y direction and ∼0.2 nm drift in x direction. The power spectrum from motion corrected aligned average clearly showed the gold ⟨111⟩ (2.35 Å) and ⟨200⟩ (2.04 Å) signal as shown in (figure 2 E and G). The same tests were done with a Gatan 626 holder with and without liquid Nitrogen (LN2) to test for holder performance.

**Figure 1:**
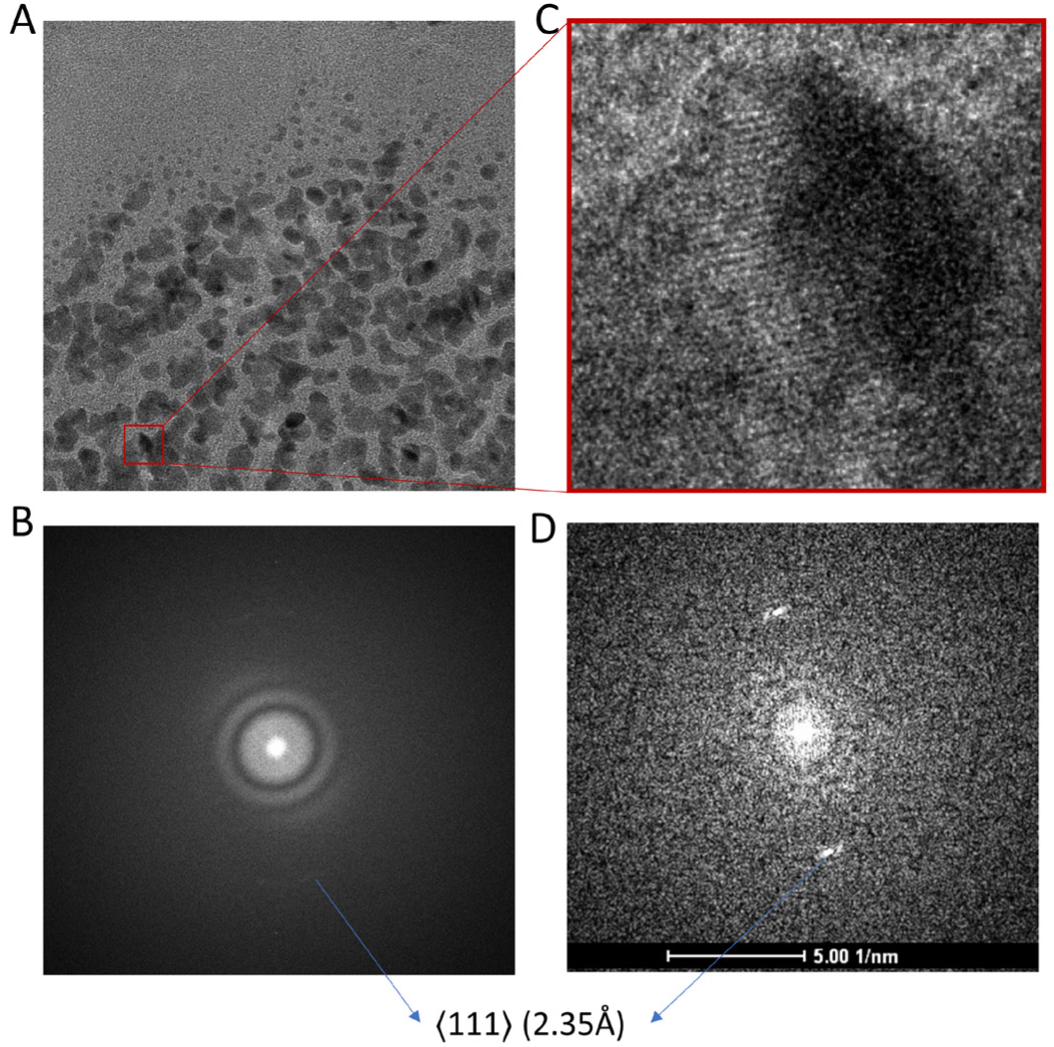
(A) Au cross grating specimen, 4 s exposure captured on BM Eagle CCD camera. (B) Power spectrum showing washout of information in one direction due to residual stage drift with weak signal corresponding to ⟨111⟩ reflection corresponding to 2.35 Å seen in the direction least affected by drift. (C) Zoomed in selected area showing Au lattice lines. (D) Power spectrum of the selected area (C) showing ⟨111⟩ reflection clearly corresponding to 2.35 Å.

**Figure 2:**
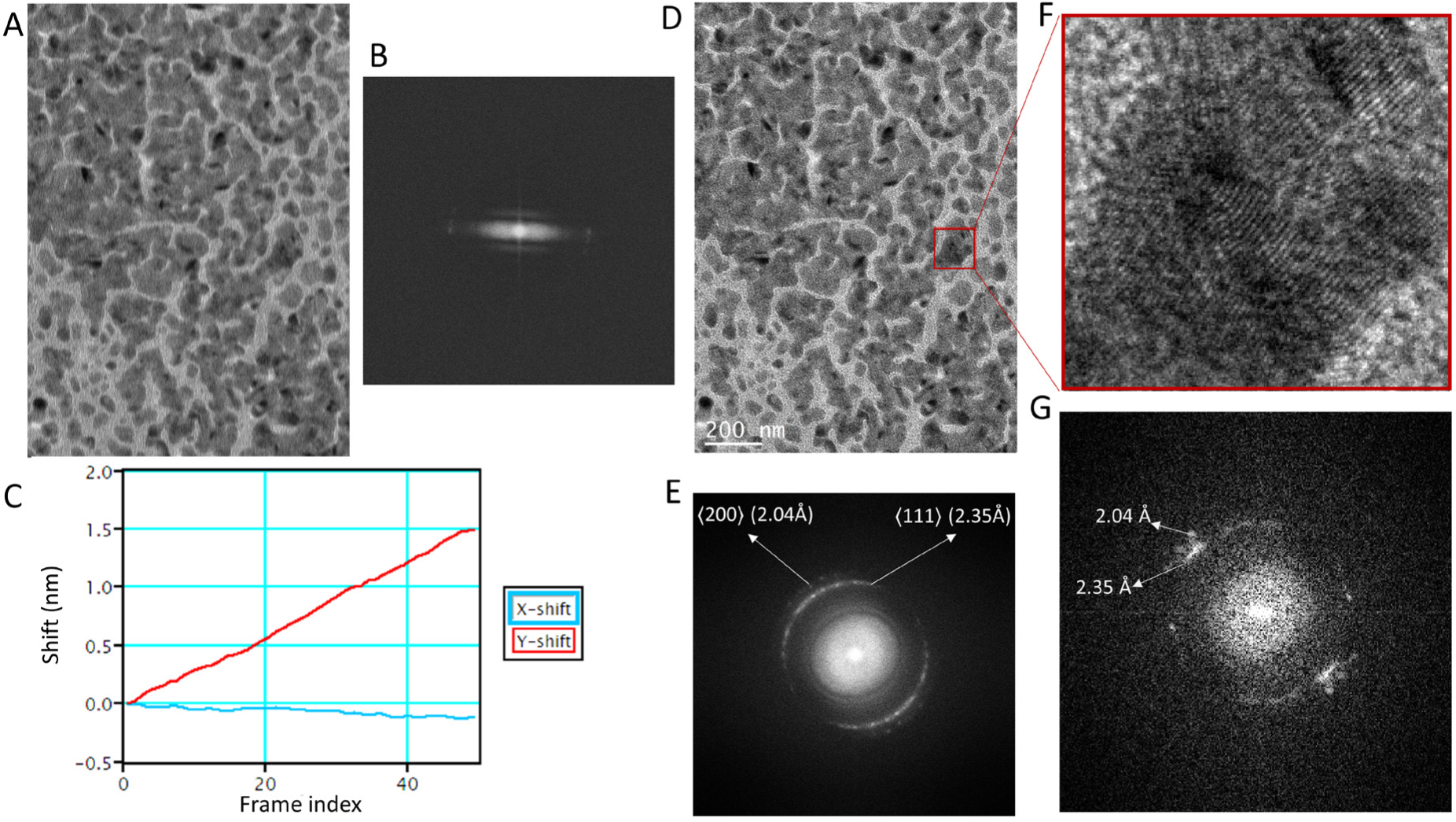
(A) Summed image of Au cross grating specimen from 60 fractions of an 8 s exposure taken using Gatan Alpine camera. (B) Power spectrum showing washout of information due to residual stage drift. (C) Motion correction in Gatan digital micrograph showing estimated motion along x and y axis. (D) Motion corrected average. (E) Retrieval of information from motion correction as shown by the power spectrum of the image in (D) showing 2.35 Å as well as 2.04 Å rings corresponding to the ⟨111⟩ and ⟨200⟩ reflections respectively. (F) zoomed in region from the original image showing Au lattice and (G) the corresponding selected area power spectrum.

In the first instance we observed wash out of Thon rings in the power spectrum caused by vibration (not fixable by drift correction). These vibrations predominantly acoustic in nature, were seen to have more pronounced impact with cryoholders likely due to the larger surface area associated with the liquid nitrogen (LN2) dewars compared to the room temperature holder (supplementary figure 1D-F). The Alpine camera power supply was identified as the reason for the acoustic vibrations and was relocated to an adjacent utility room thereby significantly attenuating these vibrations and increasing data quality (supplementary figure 1 G). Next, we proceeded to automate the usage of Alpine camera using SerialEM [30] as well as to enable automatic data collection. We performed tests with and without beam-image shift on a C-Flat grid (Protochips^TM^) to mimic single particle acquisition for collecting 9 holes per stage move. The purpose of this was to test whether switching between lower SA to high SA magnifications required for cycling between hole finding, autofocusing and data acquisition would lead to any major hysteresis in the non-constant powered TWIN objective lens. The results of CTF estimation using CTFFIND [29] showed that ∼90% of the images could be fit to a resolution of 3.5-4 Å (supplementary figure 1 G) and had an average astigmatism measure of 29.2nm (supplementary figure H). Coma-free alignment is critical for obtaining high-resolution cryo-EM datasets[30]. In contrast to higher end Thermo Fischer Scientific microscopes where coma-free alignment is accessible in the user interface, only rotation centering is available on the Tecnai G2 Spirit. This issue is easily mitigated by integrating the camera use with SerialEM[31] which makes it possible to access the tilt coils to achieve coma free alignment as well as to perform coma vs image shift calibration to reduce astigmatism and coma arising from using beam image shift. Once these calibrations were performed and sufficiently reproducible optical performance was observed, we proceeded to collect SPA datasets.

### Sub-3Å structure of Apoferritin

To assess whether the optical stability seen in a limited number of datasets measured on carbon translates to long data collections required for biological specimen, we used Apoferritin as our first test sample. We obtained a throughput of ∼170 images/hour with beam image shift data collection on 9 holes and a 15 second stage drift settling time, which was used for 4 rounds of autofocusing. For the whole dataset, the maximum resolution to which the CTF could be reliably fit using CTFFIND was Gaussian distributed around 4.5-5 Å. Further image processing yielded a 2.65 Å final structure. Although the first 20,000 particles were sufficient to reach 2.92 Å, further addition of particles did not improve the result dramatically. The calculated B-factor from Rosenthal and Henderson[32] plot was 139 Å^2^ and the sharpening B-factor from Guinier plot calculated during refinement was 110 Å^2^. Although prior to data collection the coma was corrected to a threshold of 0.1 mrad, the average residual coma measured from ctf refinement [33] for all the 9 beam image shift optics group was noted to be 0.58 and 0.60 mrad in x and y respectively.

Furthermore, analysis of Bayesian polishing[34] B-factor (figure 4D) showed steep rise of estimated B-factor as a function of dose. We compared this to polishing B-factor obtained for a megapixel equalised zero loss filtered dataset of apoferritin acquired on a S-FEG 300-keV Titan Krios captured using a GATAN K3 detector at a pixel size of 0.82 Å. At 120-keV the low-resolution contrast for non-dose weighted averages was seen to drop progressively with every 10 e^−^ Å^−2^ accumulated dose. The low-resolution contrast for the 300-keV dataset albeit being poor when compared with corresponding dose fractions from the 120-keV dataset, followed the same pattern of attenuation with the progression of dose (Figure 4A). However, for high-resolution information, the resultant reconstructions showed that in the 120-keV dataset compared to the 300-keV dataset very little high-resolution information is present post 30 e^−^ Å^−2^(Figure 4 B and C). These observations are in line with previous studies comparing 100-keV and 300-keV dose dependent loss of high-resolution signal in the diffraction pattern from 2D crystals of C_44_H_90_ paraffin and purple membrane [20].

**Figure 3:**
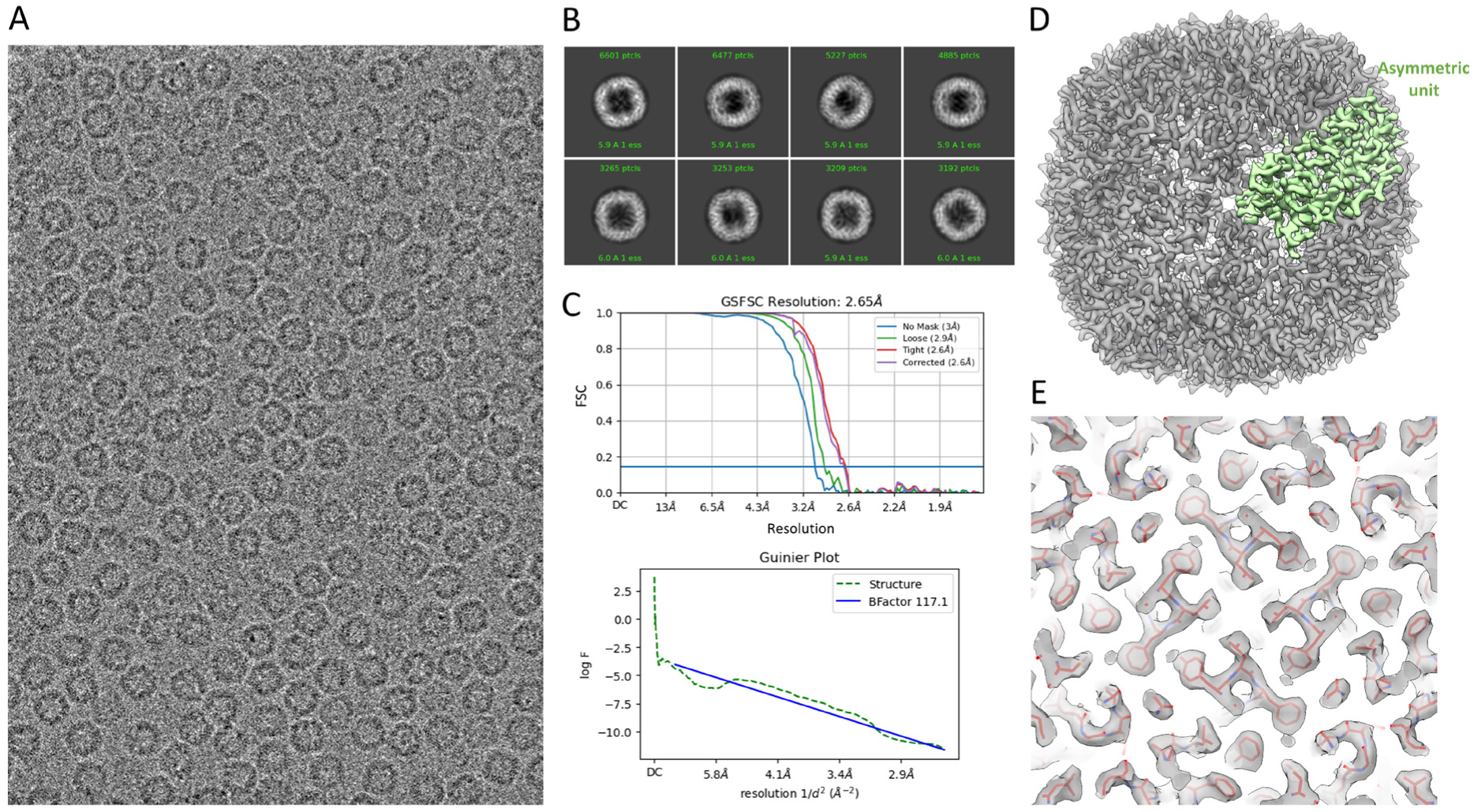
(A) Representative image of apoferritin at 0.865 μm defocus. (B) 2D class averages showing secondary structure detail. (C) FSC plot showing Gold standard resolution at 0.143 at 2.65 Å and Guinier plot showing b-factor calculated during refinement in cryoSPARC. (D) 3D reconstruction of apoferritin at 2.65 Å resolution (grey) and density in yellow corresponding to one asymmetric unit. (E) Map quality showing well fitted backbone and sidechain density from rigid body docking of pdb id: 7A6A.

**Figure 4:**
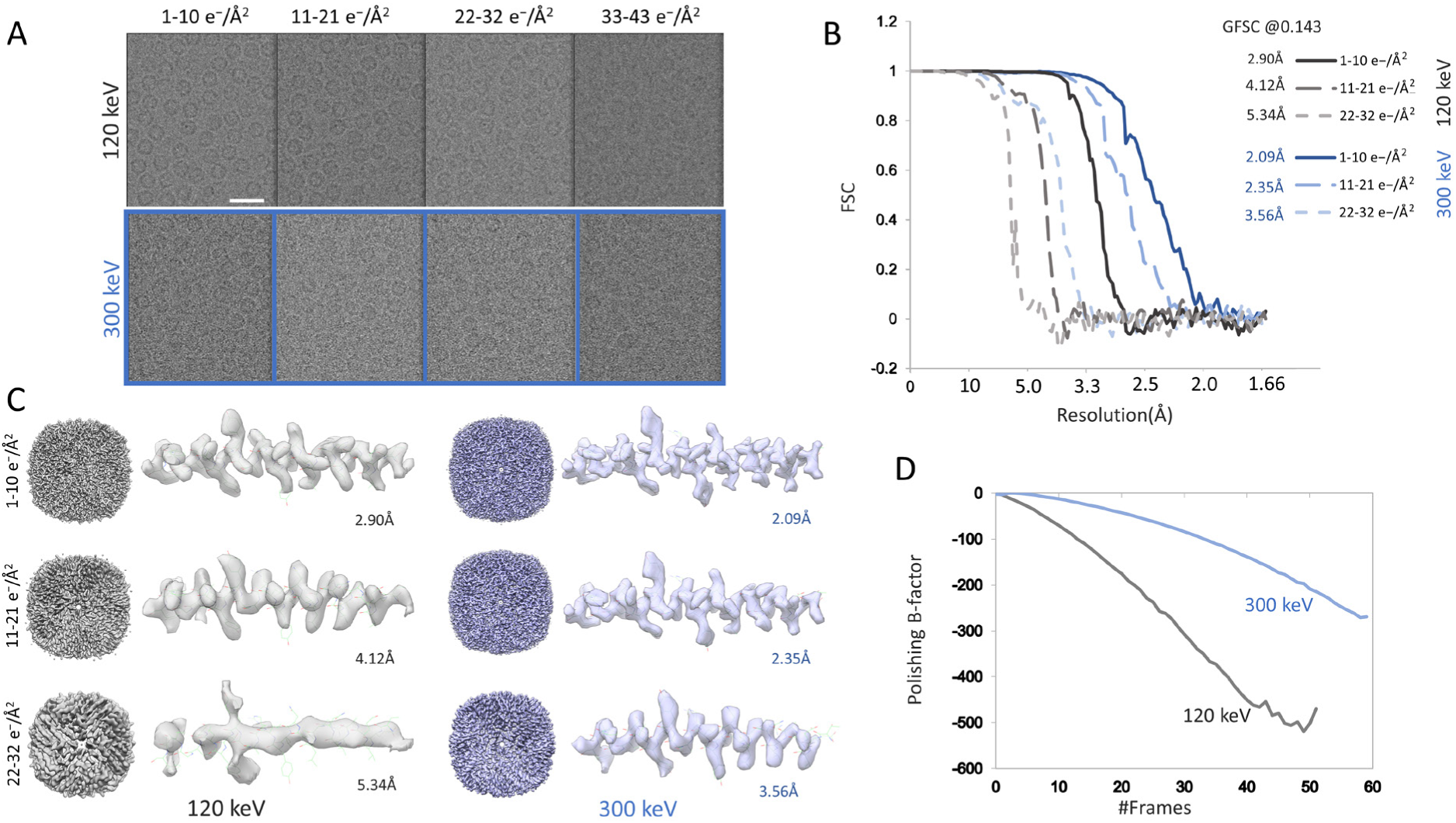
(A) Non-dose weighted, motion corrected averages corresponding to every 10 e^−^ Å^−2^, top from selected area of an apoferritin image taken at 0.865 μm defocus imaged using Tecnai G2 TWIN operated at 120-keV and captured using Gatan Alpine camera and bottom corresponding to selected area from image taken on G1 Titan Krios operated at 300-keV and captured using Gatan K3 at 0.947 μm defocus. (B) Gold standard FSC plot for reconstructions from particles for dose range until 32 e^−^ Å^−2^ for both 120-keV (grey) and 300-keV (blue). (C) 3D reconstruction for dose range corresponding to the first 32 ^−^ Å^−2^e for both 120-keV (grey and 300-keV (blue) and zoomed in region of the map showing alpha helix comprising residues 38 Phe to 11 Thr. (D) Per frame B-factor calculated during Bayesian Polishing in RELION-5 for both 120-keV (grey) and 300-keV(blue).

### High-resolution structure of a 153 kDa asymmetric dynamic membrane protein

We further tested the imaging configuration to see how it fares with a more challenging asymmetric sample and its feasibility for screening. For this purpose, we chose the 153 kDa M4 muscarinic acid receptor (M_4_ mAChR) bound to G_i1_ protein and in complex with small molecule iperoxo as well as LY298 (M4R-G_i1_-lpx-LY298), which belongs to the group of G protein-coupled receptors (GPCRs). This complex was previously reconstructed to a resolution of 2.4Å [35] from ∼6,000 zero loss filtered movies collected at 300-keV on a Titan Krios with K3 detector in CDS mode. Using the Alpine on the Tecnai G2 Spirit we collected a megapixel equalised dataset to match a typical screening dataset that we normally collect on a Talos Arctica. In ∼11 hrs we obtained ∼1,300 movies. Initial analysis using a combination of RELION-4.0 and CryoSPARC ab-initio classification followed by non-uniform refinement yielded a resolution of 5.8 Å. Further 3d classification in RELION-5.0 with BLUSH regularisation[36] followed by refinement using BLUSH regularisation yielded a final 50,031 particles that reached a resolution of 4.4 Å (figure 5). Local resolution estimates (supplementary figure 2F) showed that the resolution in the transmembrane helices (TMs) is of the order of 4.2-5 Å with highest resolution of the order of 4.2-4.8 Å for TM3. This resolution in the TMs were sufficient to unambiguously trace the backbone as shown by the rigid body fit of the previously published M4R-G_i1_-lpx-LY298 structure (supplementary figure 2A). The resolution was sufficient to unambiguously identify a significant tilt in the receptor with respect to the G protein when compared to that solved previously (supplementary figure 2E). The structure had sufficient resolution to identify the presence of the small molecule (LY298) occupancy in the allosteric binding site whereas the orthosteric inhibitor iperoxo was not visible (supplementary figure 2D). Notably, in the previously published reconstruction from the 300-keV dataset [35] a density for iperoxo was detectable, but the alkyne bond was not visible. Since the 300-keV dataset had 617,793 particles compared to the 50,031 particles in the current study, it is difficult to ascertain whether the lack of the iperoxo density in the structure is due to a poorer S/N ratio of individual particles compared to the zero loss filtered 300-keV FEG dataset or is a result of partial occupancy or dynamics which would have benefited from averaging more particles. Although the resolution is not sufficient to unambiguously confirm the identity of LY298, the results show that this resolution level is sufficient to identify the presence or absence of a small ligand at the binding site.

**Figure 5:**
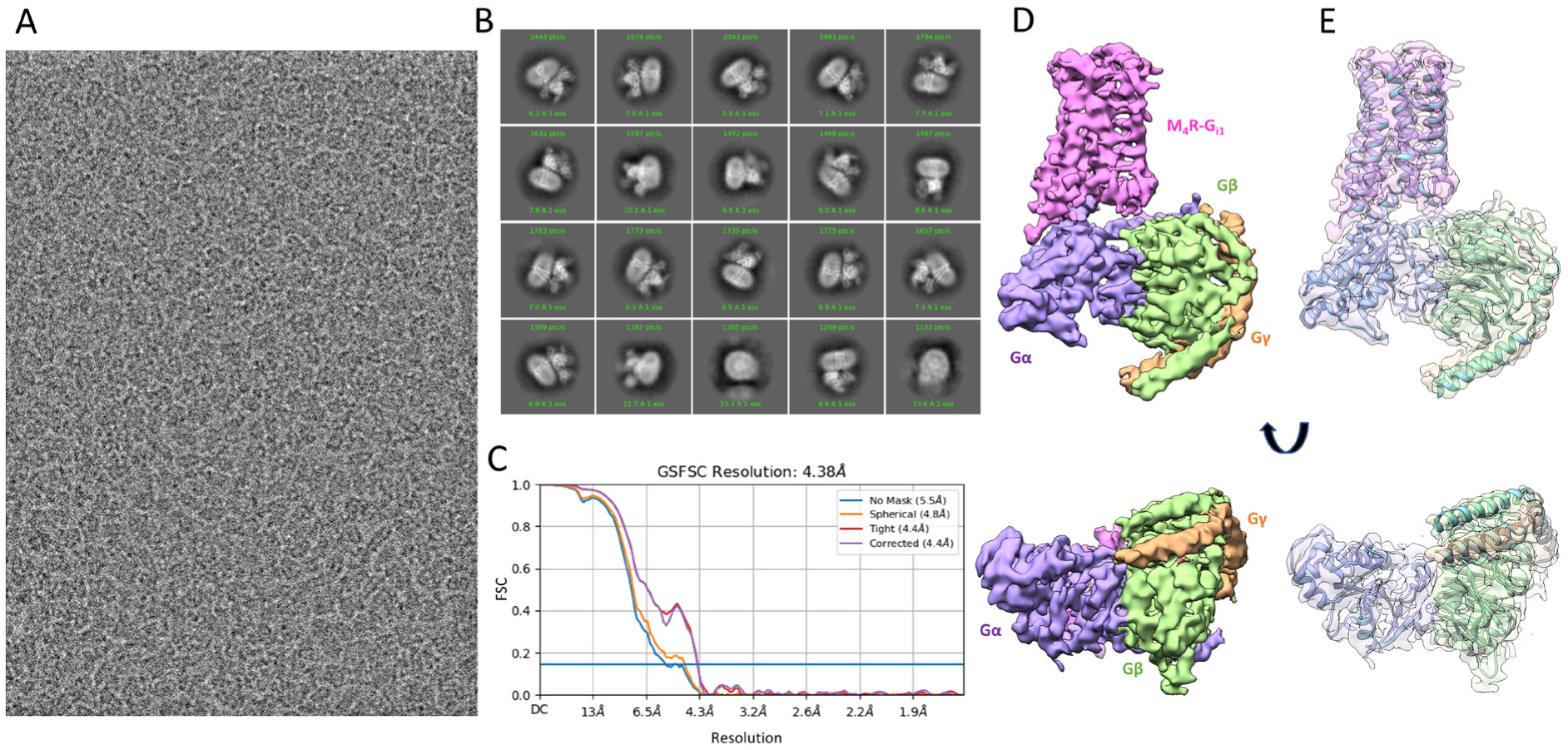
(A) Representative image of M4R-G_i1_-lpx-LY298 at 0.819 μm defocus. (B) 2D class averages showing secondary structure detail. (C) FSC plot showing Gold standard resolution at 0.143 of 4.4 Å. (D) 3D reconstruction of M4R-G_i1_-lpx-LY298 at 4.4 Å resolution showing the trans membrane helices of the receptor (purple) and the attached intracellular G-protein complex stabilised by scFv16(masked out). (E) Showing pdb id: 7TRP rigid body docked into the cryo-EM reconstruction.

### A 4.33 Å structure of 64 kDa protein haemoglobin

We further characterised the setup for imaging protein with molecular mass smaller than. 100 kDa. For this purpose, we chose human haemoglobin, a ∼64-kDa heterotetrametric heme-containing protein. The data collection strategy was kept similar to that described for the GPCR and apoferritin samples.

Haemoglobin particle distribution was less than ideal with more particles present towards the lip of the holes. About 1,300 movies were collected between two different grids. Movies where the maximum observable resolution as reported by CTFFIND that exceeded 6 Å were discarded. This resulted in ∼700 useable movies. This dataset was sufficient to get 2D class averages with secondary structure level detail. For the 3D refinement, the conventional ab-initio classification with a 30 Å initial search resolution cut-off proved to be intractable to result in a sufficient quality ab-initio model. Instead, a initial low-resolution cut-off of 12 Å and high resolution limit of 4 Å proved to be necessary to give a robust ab-initio classification result. Particles corresponding to the resultant ab-initio class that had secondary structure detail were then further subjected to homogenous refinement to get a ∼6 Å resolution structure. This structure was used as initial model for 3D classification using BLUSH regularisation in RELION-5.0. Multiple classification rounds were required to home in on a final set of ∼8,000 particles which contributed to the final reconstruction with 4.33 Å resolution (gold standard FSC (0.143)). Local resolution estimates (supplementary fig 3D) showed that outer helices in both the alpha and beta chains were exhibiting lowest resolution of ∼5.3 Å whereas the highest resolution of 4.0 Å was seen for the inner helices (supplementary fig 3 A and D). The density of heme and coordination with His-87 was also clearly visible as seen by the rigid body docked pdb id 5NI1 to the EM density (supplementary fig 3 B and C).

## Discussion

Our experiments show that a ubiquitously available 120-keV LaB6 electron microscope such as a Tecnai G2 Spirit TWIN can be retrofitted with a sub 200-keV optimised direct electron detector to massively boost its performance level. The cost associated with procuring as well as maintaining these standard 120-keV microscopes is much lower than a modern autoloader system like a Thermo Fischer Scientific Talos Arctica, Glacios, Jeol CRYO ARM 200 or even FEG based side entry holder driven microscopes such as the Talos F200X or Jeol JEM-F200. We were able to successfully integrate the Alpine detector and make it amenable for automated data collection through SerialEM and achieve a working throughput of roughly 170 movies/hr in CDS mode. The 2.65Å structure of apoferritin achieved using this setup shows that with proper sample preparation the datasets have enough optical quality to reach a resolution where model building is possible. The results from Bayesian polishing routine from RELION [34] showed that the polishing B-factor increases more steeply for the 120-keV data set compared to the 300-keV dataset (figure 4D). We do note here that the 300keV dataset was collected using zero-loss filtering, which is known to increase S/N ratio across resolution spectrum [37]. The progressive reduction in high-resolution signal per frame in the 120-keV dataset is likely due to the beam damage caused from the lower keV electrons combined with steep CTF envelope owing to poorer coherence of the LaB6 source. Since the lower keV images have very good low-resolution contrast but very little high-resolution information available past 30 e^−^ Å^−2^, the typically used total dose of 50-60 e^−^ Å^−2^ for 200-300-keV imaging may not be required for 120-keV data collection. The lower dose requirement would further shorten the exposure time required to 3-4 s allowing for increase in throughput. This would be significantly beneficial for challenging asymmetric or flexible targets where increasing the number of particles available for characterisation becomes important. These observations are consistent with previous studies [20, 21] where it has been established that 100-keV electrons have more signal producing elastic scattering as opposed to 300-keV for a given dose. Collecting more data with shorter exposure times at a lower dose, is hence a better data collection strategy for imaging at 120-keV. Although the non-CDS mode imaging is not tested in the present study, once employed should further improve throughput.

The ability of the present imaging configuration to get to a resolution of 4.4 Å for challenging samples like the 153 kDa GPCR (M4R-G_i1_-lpx-LY298) as well as its ability to successfully achieve a 4.33 Å reconstruction for even smaller 64 kDa protein target like the C2 symmetric human haemoglobin complex, with sparse data, shows the potential of this configuration to achieve below 3-4 Å resolution by collecting larger datasets. The resolution achieved in both cases were enough to ascertain the presence or absence of ligands within the complex especially as seen with the Haem complex in haemoglobin molecule obtained with only 8,865 particles (supplementary figure 3). The latest developments in motion correction algorithm in MotionCor3 as well as RELION-5’s BLUSH regularisation technique was critical for 3D classification in order to glean more information from these sparse datasets. Furthermore, the contrast available at low defocus makes imaging at lower keV an ideal candidate for imaging low molecular weight protein targets as shown in supplementary figure 4. There is sufficient low-resolution contrast to visually identify particles even at defocus as low as ∼0.6 μm for GPCR as well as haemoglobin. The above results show that this imaging configuration is an effective SPA screening tool and if needed, with larger datasets, can be used to get to a resolution range of 3Å where one can confidently do model building. Samples such as GPCRs, which require screening and characterising not just on the grid-square level variations but also hole to hole level variations, greatly benefit from screening on TEMs with multi sample loading system. Development of multigrid side entry cryo-capable holders which can accommodate 3-4 grids would be game changing for using existing 120-keV microscopes for screening purposes. Automated refilling or longer holding times for cryo-conditions to be maintained overnight without the need for manual refill would also allow for longer data collection.

In summary, widely available low-cost 120-keV LaB6 microscopes once enabled with affordable sub 200-keV direct electron detector like GATAN Alpine detector can massively bring down the economic entry barrier of cryo-EM. This could truly democratise cryo-EM with the biggest impact likely on the widespread accessibility of sample screening and optimisation.

## Methods

Alpine detector was bottom mounted as shown in (supplementary figure 1 A&B). The camera was allowed to access the Tecnai G2 Spirit TWIN pre specimen blanker (gun deflector) via a direct connection from TEM lens control electronics to the Gatan camera controller. This allowed the camera to interact with the microscope as a standalone camera. Further integration of the camera with the microscope was done through SerialEM[31] (Ver: 4.0.28). Standard serial EM calibration were done in order to efficiently use the camera along with the microscope.

### Data collection

Tecnai G2 Spirit TWIN (Thermo Fischer Scientific) equipped with a standard LaB6 thermionic emission sources (DENKA LaB6, 15 µm radius microflat surface with 90° cone angle), was operated at 120-keV. A C2 aperture of 50 µm was chosen and microscope was operated in nanoprobe mode. A nominal magnification of 60,700x at camera level corresponding to a calibrated pixel size of 0.4055 Å (super resolution mode) was chosen for data collection and beam was set to satisfy parallel illumination. The camera was operated in CDS mode. All resultant movies were Fourier binned(2X), dose weighted and motion-corrected using UCSF Motioncor2 (version 1.6.3)[7] to output both dose-weighted and non-dose-weighted average with a resultant pixel size of 0.811 Å. The non-dose-weighted averages were used for estimating contrast transfer function (CTF) parameters using CTFFIND 4.1.8[29], using RELION 4.0.1 [38] wrapper.

### Holder and optics stability measurements

Movies were recorded with at a dose rate of ∼6.5 e^−^ pixel^−1^ s^−1^. Total exposure time of 7s was used to accumulate a typical SPA dose of 60.4 e^−^ Å^−2^. Automatic data collection using serial EM was performed to collect a series of test images. Low magnification map was collected on a relatively flat region of an Au cross grating (TED PELLA, INC). Points for image acquisition were set 1-2 µm apart for automated data collection to mimic SPA style data collection routine. A drift settling time of 15 s spent doing autofocusing was used to minimise residual stage drift prior to data collection. On axis data collection as well as beam image shift data collection using 9 acquisition per stage shift pattern with and without coma versus image shift compensation were performed to test for instabilities in imaging that might arise from optics, environmental or holder instabilities. Data was collected on the same Au cross grating grid mounted on room temperature holder and Gatan 626 holder at room temperature as well as at −172 °C. Prior to high-resolution data collection, the liquid Nitrogen (LN2) level were ensured to be below the temperature transfer rod in the cryoholder. Periodic LN2 filling of the cryoholder was performed as required ensuring the above conditions to be met.

### Single particle data collection

Prior to imaging all cryo-EM samples, the astigmatism and coma was corrected as well as coma vs image shift calibrations were done using SerialEM on a C-Flat grid (Protochips) mounted on room temperature holder. Once the beam settings were transferred to SerialEM for acquisition and autofocus setting, the room temperature holder was removed and the cryo-EM grid was mounted on the stage using a Gatan 626 holder. The filament was kept running during the process.

### Apoferritin

#### Grid preparation

3µl of purified apoferritin from Thermo Fischer Scientific (VitroEase™ Apoferritin Standard) at a concentration of 4mg/ml was applied on UltraAuFoil [39] R 1.2/1.3 300 mesh. The grids were glow discharged for 30 s at 30mA in atmosphere prior to application of the sample. The excess protein solution was blotted off using a blot force of −1 and blot time of 3 seconds in a Mark IV vitrobot (Thermo Fischer Scientific) at 100% humidity and 4°C.

#### Data collection

Beam-image shift data collection with 9 holes acquisition per stage shift was performed with coma compensation on. Movies were collected with a total dose of 50.3 e^−^ Å^−2^ accumulated over 6.5 s exposure time at a dose rate of 5.25 e^−^ pixel^−1^ s^−1^ fractionated into 52 frames. A total of 923 movies were collected in a time frame of 5.45 hrs.

#### Image processing

Particle picking was performed using Gautomatch(0.53) (https://www.mrc-lmb.cam.ac.uk/kzhang/) on first 100 images and extracted binned 4 times using RELION-4.0.1. These particles were then subjected to 2D classification followed by ab-initio and 3d refinement in cryoSPARC 4.2.0 [40]. The coordinates were then exported back to RELION-4.0.1 using pyem v0.5 and re-extracted in RELION-4.0.1 centred on refined coordinates. The particles were then used for training using Topaz picker[41] and the trained model was used for particle picking on the full dataset. The resultant particles were then extracted binned 4 times and were imported to cryoSPARC 4.2.0 [40]. After 2 D classification and 3D homogenous refinement the final set of particles (127,118) were arrived upon (Extended data table 1). This final set of particles were then re extracted similarly as described above at native pixel size and were further processed in cryoSPARC 4.2.0 [40]. The refined particles were then subjected to Bayesian Polishing in RELION-4.0.1 and were then further processed in cryoSPARC 4.2.0 [40]. Post several rounds of heterogenous refinement to exclude noisy particles and CTF refinement yielded 2.65Å map (gold standard FSC 0.143 criteria)

#### Dose damage estimation

300-keV data collection: Apoferritin was frozen as mentioned above and data was collected using a G1 Krios (Thermo Fischer Scientific) operated at 300-keV, C2 aperture of 50 µm, energy filtered TEM mode with imaging done on a Gatan K3 direct electron detector equipped post a Gatan BioQuantum energy filter. Imaging was performed at a nominal magnification of 105,000X, with zero loss filtering done using a 10eV slit width. The K3 was operated in CDS mode with an effective pixel size of 0.82 Å. A dose rate of 8.9 e^−^ pixel^−1^ s^−1^ was used to accumulate a total dose of 60.21 e^−^ Å^−2^ over 5.0 s exposure time, this dose was further fractionated into 60 frames. Automatic data collection using beam image shift was performed using EPU software (Thermo Fischer Scientific).

Processing was done as mentioned above with the only exception that Bayesian polishing step was not performed. A final particle set of 106,568 particles resulted in 1.89 Å (gold standard FSC 0.143 criteria).

#### Image processing

Both the 120-keV as well as 300-keV final set of particles were re-extracted from non-dose weighted motion corrected averages corresponding to every 10 e^−^ Å^−2^ increment. The dose limited re-extracted particles were auto refined with their respective full dose final reconstructions as initial model. Initial low pass filtering for refinement was restricted to 12 Å. The estimated resolution from each dose limited particle set along with their corresponding FSC (gold standard 0.143 criteria) plot is shown in figure 4 B.

### M4R-G_i1_-lpx-LY298

#### Grid preparation

3ul of purified M4R-G_i1_-lpx-LY298 purified as described previously [35], was applied on UltraAuFoil R 1.2/1.3 300 mesh at a concentration of 15mg/ml. The grids were glow discharged for 180 s at 15mA in atmosphere prior to application of the sample. The excess protein solution was blotted off using a blot force of 10, blot time of 2.5 seconds in a Mark IV vitrobot (Thermo Fischer Scientific) at 100% humidity and 4°C.

#### Imaging

Beam-image shift data collection with 9 holes acquisition per stage shift was performed with coma vs image shift beam shift compensation on. Data was collected in CDS mode with a super resolution pixel size of 0.4055 Å. Movies were collected with a total dose of 60.53 e^−^ Å^−2^accumulated over 6.09 s exposure time at a dose rate of 6.74 e^−^ pixel^−1^ s^−1^ fractionated into 52 frames. A total of 1324 movies were collected in a time frame of ∼11 hrs.

#### Image processing

The resultant movies were Fourier binned(2X), dose weighted and motion-corrected using UCSF Motioncor3[7] to output both dose-weighted and non-dose-weighted averages with a resultant pixel size of 0.811 Å. Particle picking was performed using Gautomatch(0.53) on first 100 images and extracted binned 4 times. These particles were then subjected to 2D classification followed by ab-initio and 3d refinement in cryoSPARC 4.2.0. The coordinates were then exported back to RELION using pyem v0.5[42] and re-extracted in RELION-5.0 centred on refined coordinates. The particles were then used for training using Topaz picker[41] through RELION5.0 wrapper and the trained model was used for particle picking on the full dataset. This resulted in 479,634 particles which were then extracted binned 4 times and were imported to cryoSPARC 4.2.0. After 2 D classification and 3D homogenous refinement the final set of particles 75,816 were arrived upon (Extended Data Table 1). This final set of particles were then re extracted similarly as described above but with binning 2 and were further processed in cryoSPARC 4.2.0. The refined particles were then subjected to Bayesian Polishing in RELION-5.0 and were then re imported and further processed in cryoSPARC 4.2.0. Post several rounds of heterogenous refinement to exclude noisy particles and non-uniform-refinement[43] yielded 5.36 Å map (gold standard FSC 0.143 criteria).

Volume erase tool in UCSF Chimera[44] was used to remove the micelle density from the map and a new mask containing only the TM region and the G-protein was prepared from this map in RELION-5.0 and was used for further 3D classification with BLUSH regularisation on full dataset. The highest resolution class corresponding to 50,089 particles and were then reextracted at native pixel size and subjected to mask refinement with BLUSH regularisation. This pushed the resolution to 4.8 Å (gold standard FSC 0.143 criteria). Polishing followed by per particle defocus refinement further improved the map to achieve 4.38 Å resolution (gold standard FSC 0.143 criteria) (figure 5 C).

### Haemoglobin

#### Protein purification

Hb was purified from human blood collected from a consenting, healthy adult volunteer in accordance with 2022-30658-70864 approved by the Monash University Human Research Ethics Committee. Erythrocytes were isolated via centrifugation and cell pellets were diluted in lysis buffer (50 mM Tris, 200 mM NaCl, pH 7.4). To isolate Hb, erythrocytes were lysed with a tight-fit dounce homogeniser and clarified by centrifugation at 18,000 rpm. Cell lysate was further purified using a Superdex 26/600 S200 size exclusion column (Cytiva) pre-equilibrated in SEC buffer (50 mM Tris, 200 mM NaCl, pH 7.4). Fractions were collected and pooled and concentrated to ∼10 mg/ml using a 30-kDa spin filter column (Millipore) before being stored at −80°C.

#### Grid preparation

3ul of purified haemoglobin sample at a concentration of 10 mg/ml was applied on UltraAuFoil R 1.2/1.3 300 mesh. The grids were glow discharged for 30 s at 30mA in atmosphere prior to application of the sample. The excess protein solution was blotted off using a blot force of −1, blot time of 3 seconds in a Mark IV vitrobot (Thermo Fischer Scientific) at 100% humidity and 4°C.

#### Imaging

Beam-image shift data collection with 9 holes acquisition per stage shift was performed with coma vs image shift beam shift compensation enabled. Data was collected in CDS mode with a super resolution pixel size of 0.4055 Å. Movies were collected with a total dose of 50.14 e^−^ Å^−2^ accumulated over 6.711 s exposure time at a dose rate of 5.07 e^−^ pixel^−1^ s^−1^ fractionated into 55 frames. A total of 1368 movies between two grids were collected.

#### Image processing

The resultant movies were Fourier binned(2X), dose weighted and motion-corrected using UCSF Motioncor3[7] to output both dose-weighted and non-dose-weighted average with a resultant pixel size of 0.811 Å. Particle picking was performed using Gautomatch (0.53) on first 100 images and extracted binned 4 times. These particles were then subjected to 2D classification followed by ab-initio and 3d refinement in cryoSPARC 4.2.0. The coordinates were then exported back to RELION using pyem v0.5 and re-extracted in RELION-5.0 centred on refined coordinates. The particles were then used for training using Topaz picker through RELION5.0 and the trained model was used for particle picking on the full dataset. This resulted in 382,710 particles. In CryoSPARC 4.2.0 ab-initio classification was performed using an initial low-resolution search cut-off of 12 Å, high-resolution limit of 4 Å and with 3 classes with class similarity set to 0. Particles corresponding to the resultant ab-initio class that had secondary structure detail were then further subjected to homogenous refinement to get ∼6 Å resolution structure. This structure was used as initial model for 3D classification using BLUSH regularisation in RELION-5.0. Multiple classification rounds were done to reach a final set of 8,865 particles which contributed to the final reconstruction with 4.33 Å resolution (gold standard FSC 0.143) (figure 6 C).

**Figure 6:**
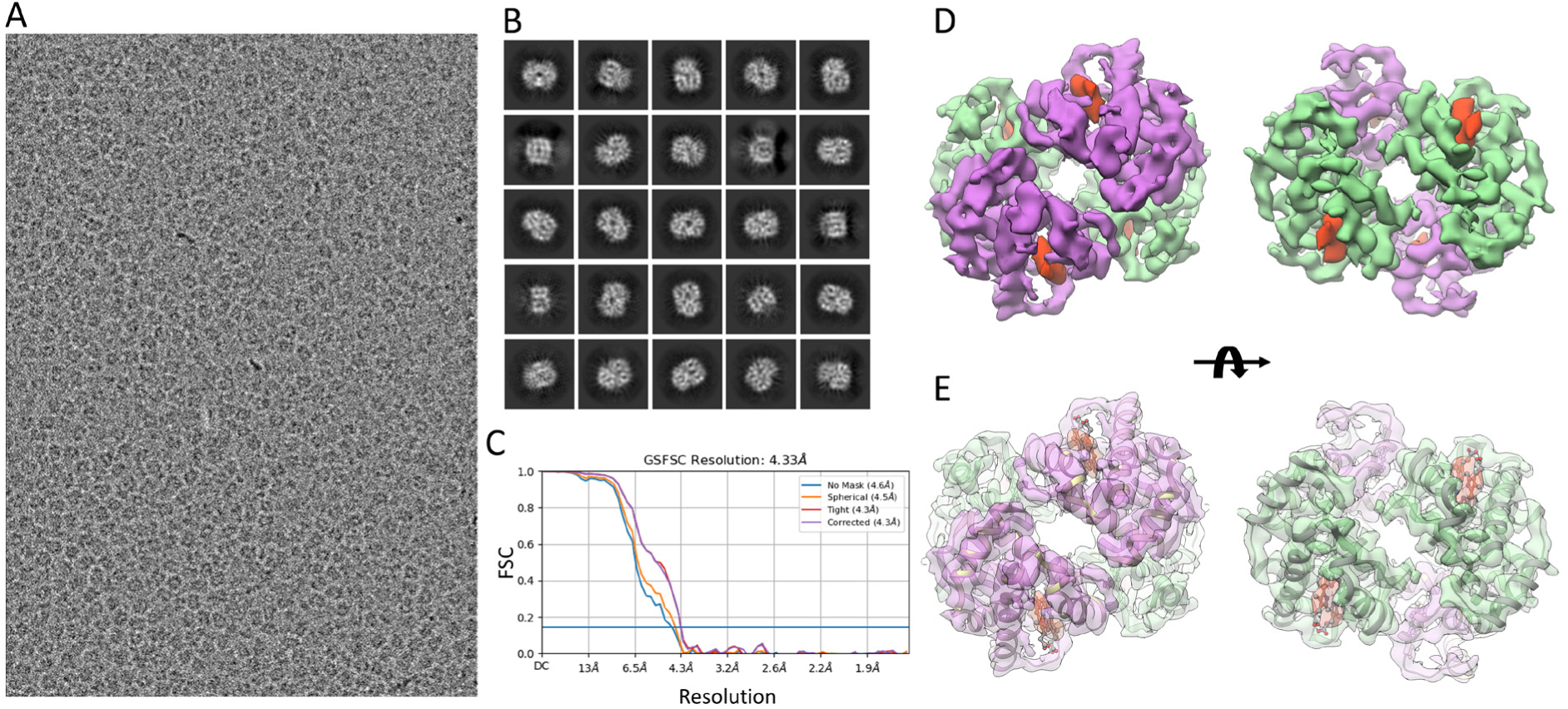
(A) Image of human haemoglobin at 0.853 μm defocus. (B) 2D class averages showing secondary structure detail. (C) Resolution according to Gold standard FSC at 0.143 of 4.33 Å. (D) C2 symmetric 3D reconstruction of human haemoglobin with purple density showing alpha chain, green showing beta chain and haem density in red and (E) showing pdb id: 5NI1 rigid body docked into the cryo-EM reconstruction.

Visualisation and rigid body docking for all structures were performed using UCSF Chimera[44] (Ver 1.16)

## Data availability

Maps have been deposited to the EMDB with EMD-43574 for apoferritin, EMD-44343 for M4R-G_i1_-lpx-LY298 and EMD-44746 for Haemoglobin. The raw movies for apoferritin have been deposited to EMPIAR(EMPIAR-11876).

## Acknowledgments

The authors acknowledge the use of instruments and assistance at the Monash Ramaciotti Centre for Cryo-Electron Microscopy, a Node of Microscopy Australia. This research used equipment funded by Australian Research Council grant ARC LIEF (LE200100045, LE120100090, LE150100132) and the Monash University MASSIVE high-performance computing facility and supercomputing resources. This work has been made possible in part by CZI grant DAF2021-225399 and grant DOI https://doi.org/10.37921/334038myxhsa from the Chan Zuckerberg Initiative DAF, an advised fund of Silicon Valley Community Foundation (funder DOI 10.13039/100014989). Grant Wu(GATAN) performed the camera installation. We also thank Mehul Patel and Brenton Cook from Thermo Fischer Scientific for their support during installation. We thank Dr.Matthew Belousoff (Monash University), Dr. Benjamin Gully (Monash University) and Dr. Sahil Gulati(GATAN) for manuscript reviews.

## Author contributions

H.V: Conceptualization, camera integration, cryo-EM grid optimisation, imaging and analysis. Z.V, J.M, C.T and D.F: protein expression and purification. H.V, D.T, G.K, R.G, S.M, C.C and G.R project management. H.V and G.R wrote the manuscript. All the authors contributed in reviewing and finalising the manuscript.

---

**Supplementary figure 1:**
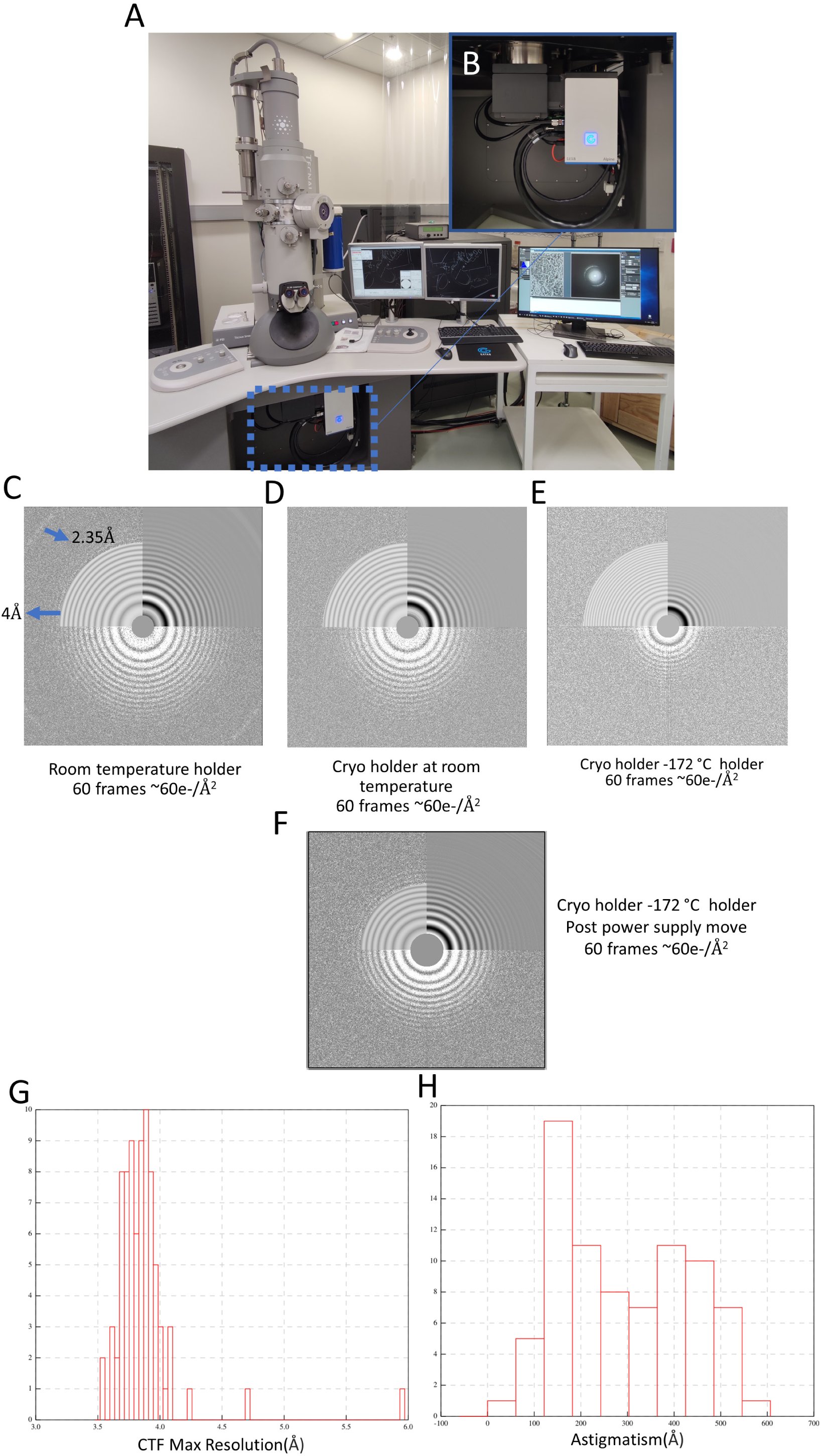
(A) Tecnai G2 TWIN 120-keV LaB6 microscope retrofitted with (B) GATAN Alpine direct electron detector. (C) Power-spectrum from Au cross grating image taken mounted on room temperature holder (right quadrant showing CTFFIND fit till 4 Å and the right quadrant with simulated fit), (D) from sample mounted on cryoholder but at room temperature showing loss of high-resolution signal (E) from image of sample mounted on cryoholder at cryo temperature showing uncorrectable vibration. (F) Power-spectrum from c-flat holey carbon grid post power supply move. (G) Astigmatism and (H) max CTF-max-resolution histogram from 80 movies collected using beam image-shift data collection.

**Supplementary figure 2:**
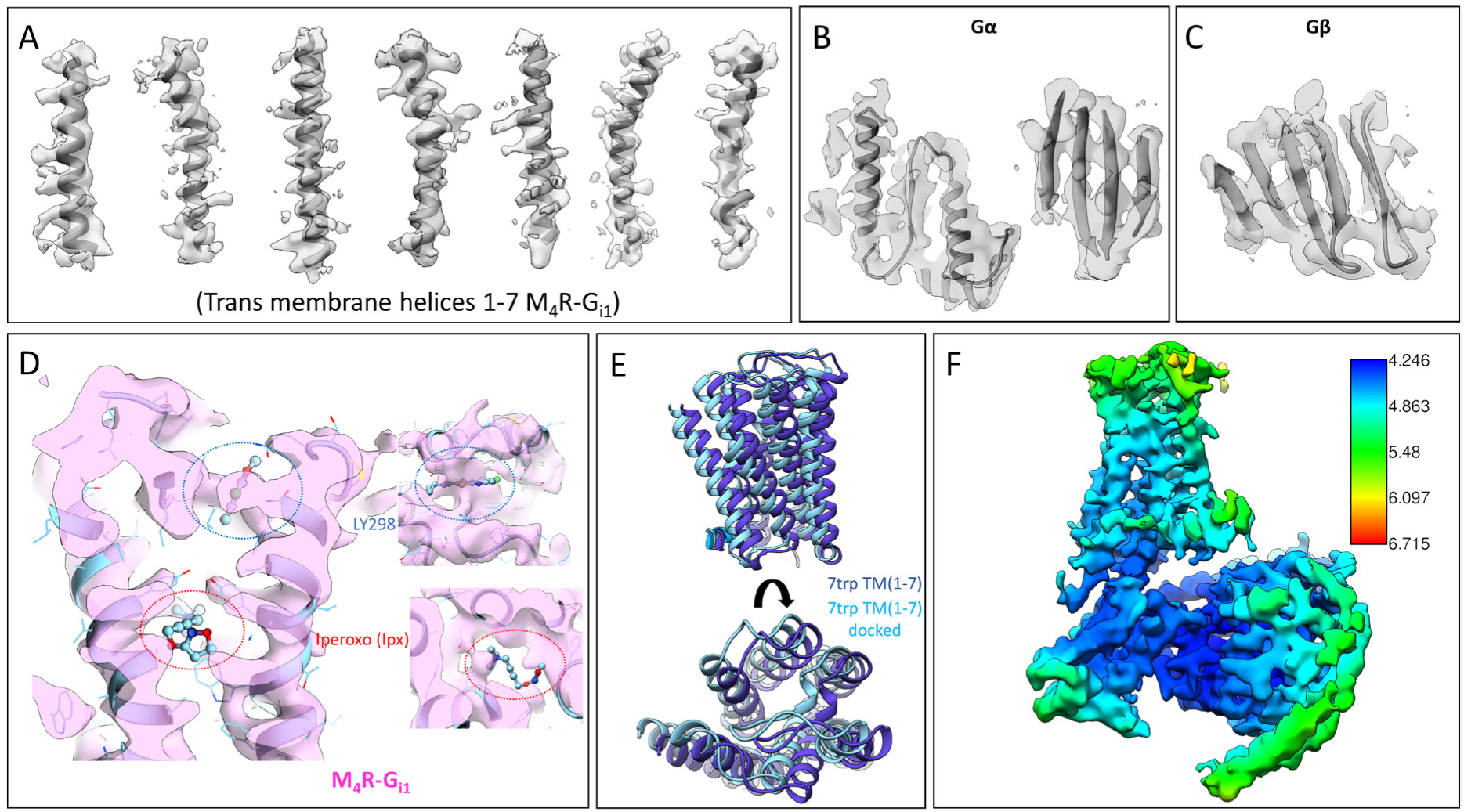
Map quality of M4R-G_i1_-Ipx-LY298 complex. (A)Cryo-EM density of transmembrane helices of M_4_R-G_i1_ rigid body docked with pdb id: 7TRP. (B) cryo-EM density from G alpha showing alpha helices with sidechain density as well as beta-sheet density. (C) Beta sheets from G beta. (D) M_4_R-G_i1_ region binding to ligand LY298 on the top density encompassed in (blue dotted line) and iperoxo missing density as shown by red dotted lines. (E) PDB id: 7TRP TMs rigid body docked to cryo-EM density and aligned with undocked pdb showing tilt with respect to the published structure. (F) Local resolution map showing differential resolution with M_4_R-G_i1_ map.

**Supplementary figure 3:**
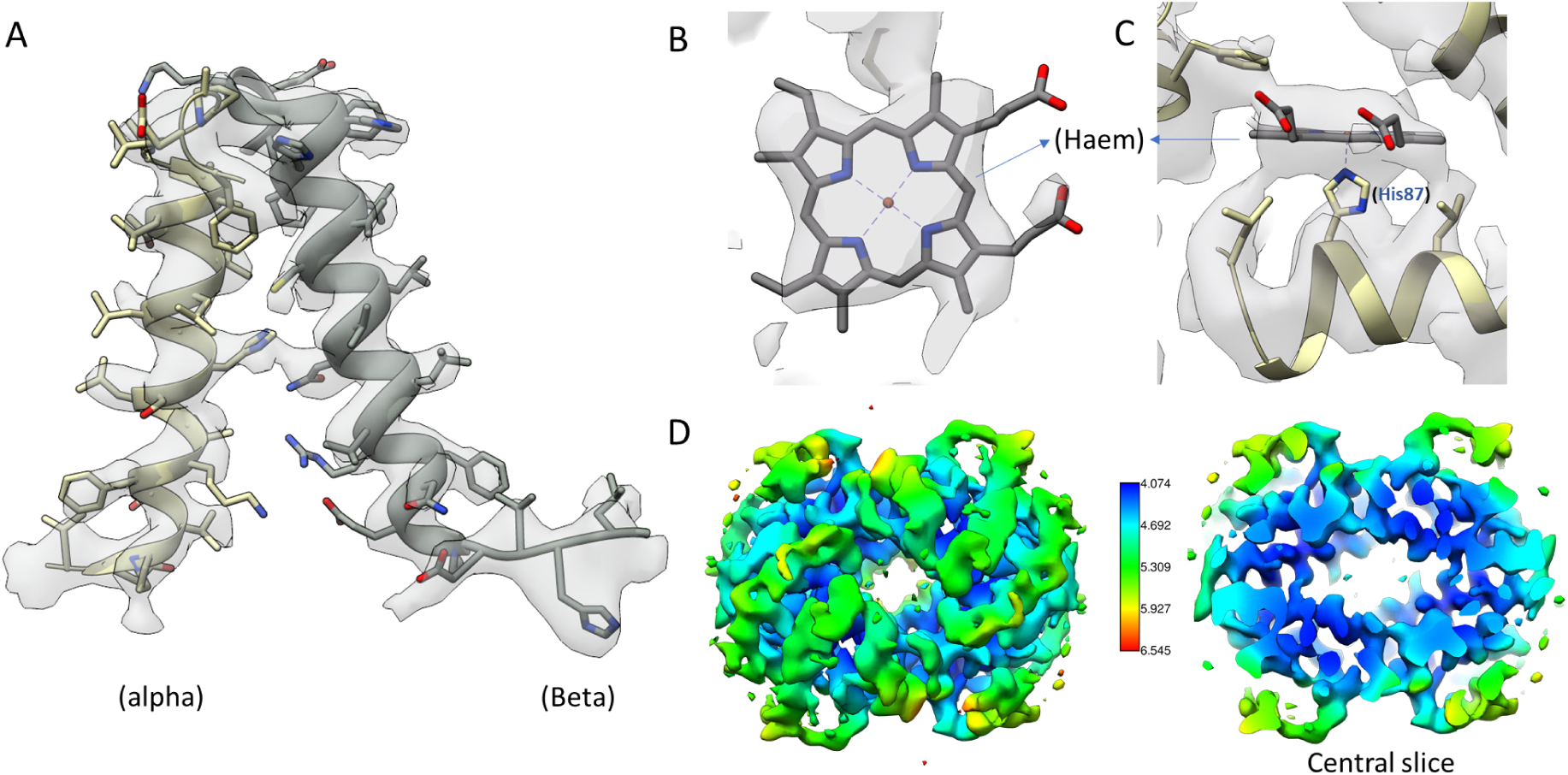
Map quality (pdb id: 5NI1 rigid body docked into the cryo-EM reconstruction): Region comprising residue Val93-His117 of alpha chain and region comprising residue His97-Glu121 of the beta chain of Haemoglobin reconstruction. (B) Haem density and (C) Density showing haem and His87 coordination (D) Map coloured according to local resolution.

**Supplementary figure 4:**
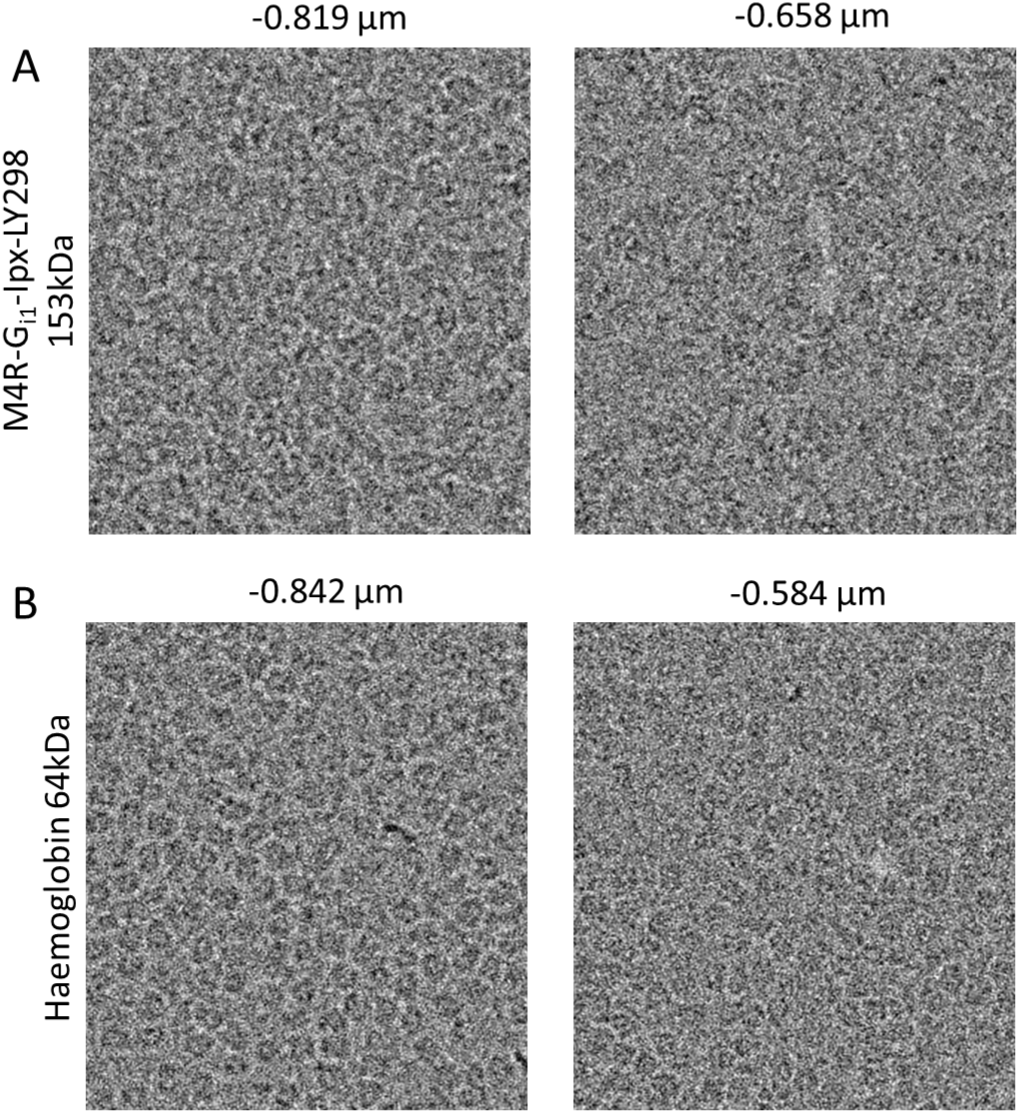
Zoomed in images of (A) M4R-G_i1_-Ipx-LY298 and (B) Haemoglobin collected on UltrAUfoil grids at different defocus (indicated above each image) showing contrast to locate particles even in images acquired with lower than a micron defocus.

**Extended Data Table 1:** Cryo-EM data collection, refinement and validation statistics.

|  | #1 Apoferritin<br>(EMDB-44745) | #2 M4R-G<sub>i1</sub>-lpx-LY298<br>(EMB-44343) | #2 Haemoglobin<br>(EMDB-44746) |
| --- | --- | --- | --- |
| **Data collection and processing** |  |  |  |
| Magnification | 60,700x | 60,700x | 60,700x |
| Voltage (kV) | 120 | 120 | 120 |
| Dose rate (e<sup>-</sup> pixel<sup>-1</sup> s<sup>-1</sup>) | 5.25 | 6.74 | 5.07 |
| Electron exposure (e<sup>-</sup> Å<sup>-2</sup>) | 50.3 | 60.4 | 55 |
| Defocus range (μm) | 0.4-0.8 | 0.5-1.0 | 0.5-1.2 |
| Super resolution pixel size (Å) | 0.4055 | 0.4055 | 0.4055 |
| Number of Movies | 923 | 1,324 | 508+860 |
| Number of Frames | 52 | 61 | 55 |
| Time taken (hrs) | 5.45 | ~11 | ~11 |
| Symmetry imposed | O | C1 | C2 |
| Initial particle images (no.) | 387,893 | 479,634 | 382,710 |
| Final particle images (no.) | 127,118 | 50,089 | 8,865 |
| Map resolution (Å) | 2.65 | 4.4 | 4.33 |
| Gold standard FSC threshold | 0.143 | 0.143 | 0.143 |
| Sharpening B-factor (Å<sup>2</sup>) | -117.1 | -151.9 | -194.9 |

## Notes

### Competing Interest Statement

SM CC are employees of GATAN, Inc. which manufacturers and markets the Alpine direct electron detector.

